# Immune dysregulation in the prostates of C57BL/6^Aire-/-^mice mirrors that seen in human benign prostatic hyperplasia

**DOI:** 10.1101/2025.08.12.669857

**Authors:** Nadia A. Lanman, Meaghan M. Broman, Harish Kothandaraman, Gregory M. Cresswell, Gada D. Awdalkreem, Dilinaer Wusiman, Andree K. Kolliegbo, Alexander P. Glaser, Brian T. Helfand, Renee E. Vickman, Jiang Yang, Simon W. Hayward, Timothy L. Ratliff

## Abstract

Benign prostatic hyperplasia (BPH) is the most common urologic disease in aging men, resulting in significant morbidity. The etiologies of BPH are unknown, though chronic prostatic inflammation is known to promote hyperplasia, fibrotic remodeling, and therapeutic resistance in BPH. BPH is highly complex and heterogeneous, presenting with varying degrees of stromal and epithelial proliferation, fibrosis, inflammation and associated lower urinary tract symptoms. This complexity presents challenges in developing models. Here, we characterize an *Aire*transcription factor-deficient non-resolving (chronic) inflammation model in a C57BL/6J background for the study of the prostatic inflammation present in BPH. This chronic inflammatory model exhibits a lack of central tolerance but retains an otherwise intact and functional immune system. C57BL/6^Aire-/-^mice were subcutaneously injected with prostate homogenate protein and Freund’s Complete Adjuvant and boosted after 10 days. After 21 and 35 days, whole prostates were collected for histology, flow cytometry, and scRNA-seq, which were then compared to human BPH scRNA-seq data Inflammation was confined to the prostate in C57BL/6^Aire-/-^mice. Histological and scRNA-seq data show that the dominant leukocyte phenotypes in the prostates of C57BL/6^Aire-/-^mice are B and T lymphocytes. Macrophages in C57BL/6^Aire-/-^mouse prostates express signatures associated with an array of phenotypes as also seen in BPH. We identify a *Trem2*^+^population of macrophages, and aging-associated Cd8^+^*GZMK*hi *GZMB*low T cells in C57BL/6^Aire-/-^prostates similar to those seen in human BPH. Further, fibroblast clusters in C57BL/6^Aire-/-^are similar to fibroblasts identified from the prostates of patients diagnosed with BPH, and these clusters also express markers associated with aging. Overall, the inflammation and predicted interactions between leukocytes and stromal cells observed in the prostates of C57BL/6^Aire-/-^mice resemble human BPH, making this model useful for studying the impact of inflammation-driven prostatic hyperplasia.

## Introduction

Benign prostatic hyperplasia (BPH) affects nearly 75% of men aged 60-69 and over 80% of men 70 and older^1^. This chronic, progressive condition involves non-malignant proliferation of epithelial and stromal cells, causing lower urinary tract symptoms like frequency, urgency, nocturia, and incomplete voiding^2^. Despite its prevalence, BPH’s underlying causes remain unknown, contributing to a 45% treatment failure rate due to incomplete understanding of pathogenesis and focus on prostate physiology rather than disease mechanisms^3^.

BPH involves complex interactions between hormonal factors^4^, epithelial-stromal interactions, genetic factors, and inflammatory pathways, causing cell growth in the periurethral transition zone^5–11^. While inflammation’s precise role in prostatic hyperplasia is unclear, recent studies suggest inflammatory signaling dysregulation promotes leukocyte accumulation and hyperplasia^9, 10,12–14^. TNF release from macrophages encourages fibroblast proliferation, while TNF blockade reduces prostatic hyperplasia in mouse models and humans and is associated with lower BPH incidence and reduced prostate size^10, 15^.

Several rodent models have been described and utilized to study prostatic inflammation and hyperplasia, including diet and hormone-inducible models^16–20^. The T+E2 hormone model mimics aging men’s hormonal environment but lacks immune specificity^18,21,22^. The *Probasin*promoter driven expression of prolactin (Pb-PRL) transgenic mouse develops prostatic hyperplasia and inflammation but has decreased androgen receptors and doesn’t respond to 5-ARIs^19,20,23,24^. NOD mice develop organ-specific inflammation including in the prostate but have a Th1 immune bias limiting population studies^25–28^. POET-3 provides antigenic specificity and demonstrates inflammation-driven hyperplasia but has limited, quickly resolving inflammatory responses^29–31^. Each model has limitations: T+E2 lacks immune specificity^18,21,22^, Pb-PRL has limited immune diversity^19,20,23^, NOD has Th1 bias^25–28^, and POET-3 has transient responses^17,29–32^. Better models are needed to mirror human BPH’s prostatic hyperplasia and non-resolving inflammation^10,14^.

This study describes novel use of the Autoimmune Regulator (Aire) knock-out mouse model (C57BL/6^Aire-/-^) for studying inflammation’s impact on prostate and hyperplasia mechanisms. Aire prevents autoimmunity by inducing deletion of high-affinity T cells in thymic epithelial cells^34,35^. Aire deficiencies cause loss of central tolerance and autoimmunity development^35,36^. The C57BL/6^Aire-/-^model exhibits chronic, non-resolving inflammation localized to the prostate (encouraged by prostate homogenate immunization as previously reported for other models^25,26^) rather than systemic inflammation, while maintaining normal immune responses^38^. This model could be valuable for studying chronic prostatic inflammation effects in human BPH. Here, we characterize the C57BL/6^Aire-/-^ mouse, study the effects of inflammation in the prostate, and compare this model with wild-type C57BL/6J mice and human BPH tissue.

## Materials and Methods

### Mice

B6.129S2-*Aire^tm^*^1^*^.1Doi^*/J (Strain 004743, The Jackson Laboratory, Bar Harbor ME) were backcrossed with C57Bl/6 mice for at least 9 generations at JAX. Aire^+/-^ mice were purchased and bred to generate C57Bl/6^Aire-/-^, C57Bl/6^Aire+/-^, and C57Bl/6^Aire+/+^ offspring. All mice were housed and maintained under pathogen-free conditions with 12 light/12 dark cycles. All mouse procedures were performed in accordance with Purdue Animal Care and Use Committee (PACUC) approved protocols.

To collect prostate homogenate protein, prostates from 8-12 week old male C57BL/6 mice were collected, minced, digested in complete RPMI + 1mg/ml collagenase for 1 hour, then heat shocked at 95°C for 5 minutes. Protein was quantified on a NanoDrop Spectophotometer (DeNovix Inc, Wilmington, DE) and stored at-20°C.

Male 8-12-week-old C57BL/6^Aire-/-^ mice were subcutaneously injected with 200μg of prostate homogenate protein and Freund’s Complete Adjuvant (Thermo Scientific, Waltham, MA) in a 1:1 ratio for a total volume of 100μl. After 10 days, mice were boosted subcutaneously with 200μg prostate homogenate protein and Freund’s Incomplete Adjuvant (Thermo Scientific) at a 1:1 ratio for a total volume of 100μl. All four lobes of the prostates were collected at 21 days or 35 days post-booster, with one of each paired lobe of every prostate fixed in 10% neutral buffered formalin (NBF) for histology and the other processed for scRNA-seq. Prostate lobes were pooled from three mice for each of the final scRNA-seq samples. Lung, kidney, liver, spleen, and pancreas were also collected and fixed in 10% NBF for histology.

### Cell flow cytometry

All four lobes of mouse prostates were collected from C57BL/6^Aire-/-^ mice, digested, and processed to a single-cell suspension as described^30,39^. Prostates were minced, digested in 1 mg/mL collagenase (Sigma-Aldrich, St. Louis, MO) in RPMI-1640 (Gibco, Waltham, MA) media containing 10% FBS (Corning, Corning, NY) with shaking at 37°C for 2 hours, then filtered through a 70μm mesh filter. Cells were suspended in 100μl complete RPMI with 2μl mouse TruStain FcX antibody (Biolegend, San Diego, CA) and 1μl Zombie Violet Fixable Viability dye (Biolegend) per sample and incubated at room temperature (RT) for 10 minutes in the dark. Cells were then divided and incubated with cocktails of cell surface antibodies (Biolegend) listed below at 1μl in 100μl PBS at 4°C for 30 minutes in the dark. Cells were rinsed and fixed in 10% NBF for 10 minutes. Samples were run on a BD Fortessa Flow Cytometer (BD Biosciences, Franklin Lakes, NJ) and analyzed with FlowJo software (BD Biosciences).

### Single-Cell RNA Sequencing

Six male C57BL/6^Aire-/-^ mice were used in scRNA-seq experiments, with prostates pooled from three mice for each scRNA-seq sample. Half of all four prostate lobes of prostates were extracted, pooled, and then digested in complete RPMI with 1mg/ml Collagenase I on a shaker at 225 RPM and 37°C for 1.5 hours. Samples were filtered through a 70μm strainer, centrifuged at 250g for 5 minutes and resuspended. Dead cells were removed using the EasySep Dead Cell (Annexin V) Removal kit (STEMCELL Technologies, Vancouver, Canada) per the manufacturer’s protocol. Cells were counted with trypan blue and diluted to a concentration of 1000 cells/μl, prepped with the Chromium Next GEM Single Cell 3’ Kit v3.1 (10x Genomics, Pleasanton, CA) per manufacturer’s protocol, and two wells per sample were loaded into the 10x Chromium (10x Genomics) instrument per the manufacturer’s protocol for a recovery target of 10,000 cells. RNA clean-up and library preparation were performed per the 10x Genomics protocols. Quality control and alignment of the single-cell RNA-seq datasets to the GRCm38 reference genome was performed using CellRanger v2.1.0 (10x Genomics, Pleasanton, CA).

Matrices were imported into R v4.1.1 and analyzed using Seurat v4.1.0 R^40,41^. Initial cell type identities were assigned using the SingleR v1.4.0 algorithm^42^ using the Immunological Genome Project reference.

## Results

### Immunized C57BL/6^Aire-/-^ mice consistently develop persistent and localized prostatic inflammation

scRNA-Seq and histologic analyses of human BPH specimens indicate that T cells are the dominant immune cell subtype, followed by macrophages^12^. Histologically, most immune cells are present in the stroma of human BPH prostates (**Fig 1A**). Stromal lymphoid cells are arranged in scattered individual cells, periglandular or perivascular loose aggregates, or organizing follicle-like structures, occasionally forming germinal center-like areas (**Fig 1A,B)**. Aggregates and organizing lymphoid structures are composed of CD4^+^ T cells mixed with CD8^+^ T cells and CD79a^+^ B cells (**Fig 1B**). Intraepithelial lymphocytes are predominately CD8^+^ T cells (**Fig 1B**). Myeloid cells, predominately macrophages and mast cells, are observed within the glandular lumina and scattered within the stroma (**Fig 1C)**.

**Figure 1.**
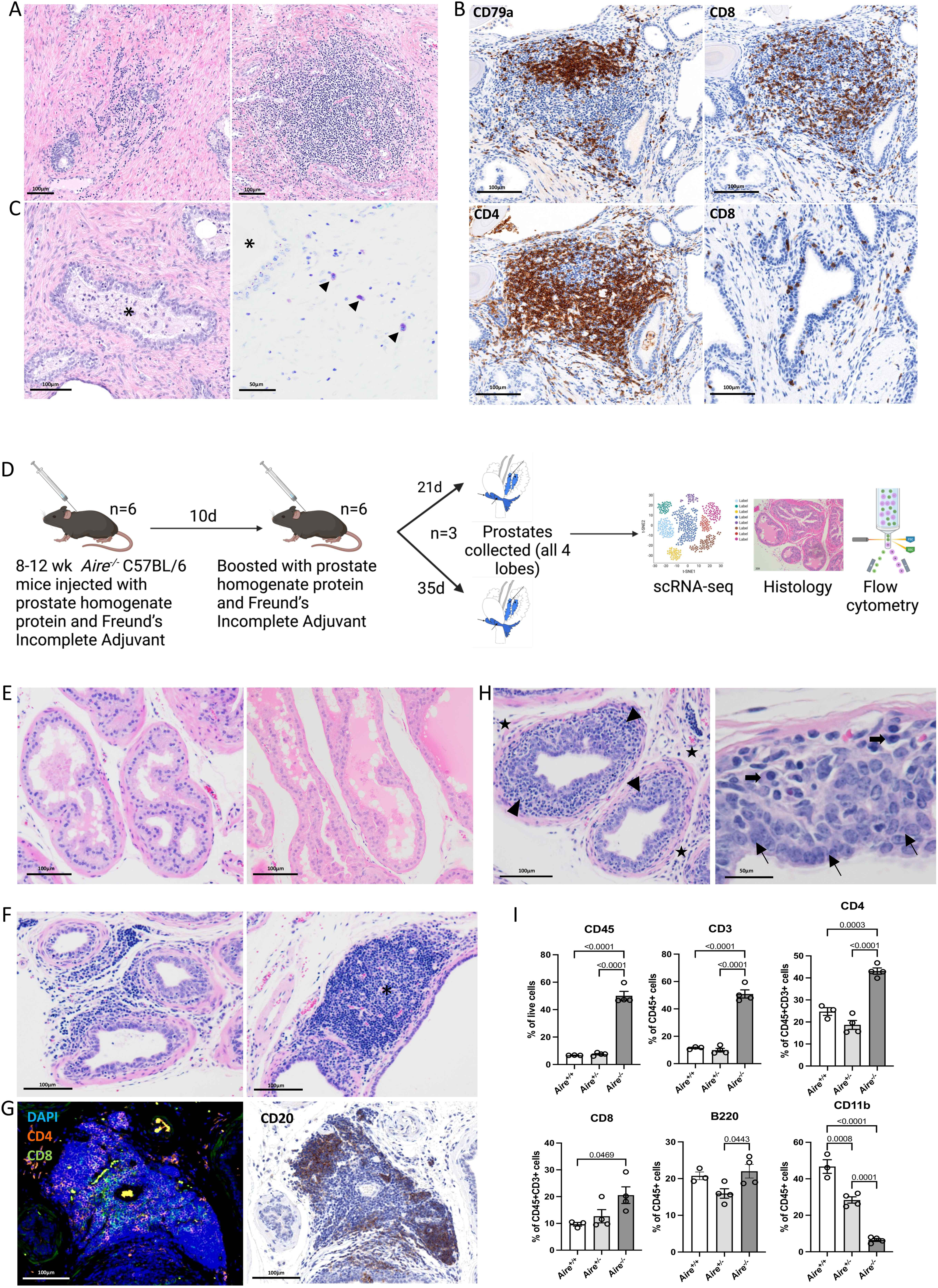
Cellular composition and architecture of C57BL/6^Aire-/-^ prostates and comparison with human prostates. A. H&E sections of human prostate immune cells showing stromal lymphoid cells in loose aggregates (left) to dense lymphoid structures (right). B. IHC of stromal lymphocytes in organizing lymphoid structures highlighting CD79a^+^ B cells (top left), CD8^+^ T cells (top right), CD4^+^ T cells (bottom left), and intraepithelial CD8^+^ T cells (arrowheads). C. H&E section (left) of human prostate showing intraluminal macrophages (asterisk). Toluidine blue-stained section (right) highlighting stromal mast cells (arrowheads) adjacent to gland (asterisk) D. An overview of the mouse experimental design. Aire knockout mice in a C57BL/6 background were injected with prostate homogenate protein and Freund’s Incomplete Adjuvant. After 10 days, mice again received a booster of prostate homogenate protein and Freund’s Incomplete Adjuvant. All four lobes of the prostates were harvested at either 21 days or at 35 days after receiving the booster and were subjected to scRNA-seq, histological analysis, or cell flow cytometry. E. H&E sections from dorsal prostate of C57BL/6^Aire+/+^ (left) and C57BL/6^Aire+/-^ (right) mice. F. H&E sections of dorsal prostate from immunized C57BL/6^Aire-/-^ mouse prostates with prominent lymphoid aggregates (left) and organizing lymphoid structures (right) with germinal center-like areas (asterisks). G. Immunofluorescence (left) and immunohistochemistry (right) highlighting arrangement of CD4 (orange) and CD8 (green) T cells and CD20 (DAB) B cells in stromal lymphoid aggregates. H. Dorsal prostate of immunized C57BL/6^Aire-/-^ mouse showing infiltration of lymphoid cells (arrowheads) adjacent to the glandular epithelium. Infiltrating cells consist of lymphoid cells including numerous plasma cells (bold arrows). Note expansion of stromal (stars) and epithelial (thin arrows) compartments compared to C57BL/6^Aire+/+^ and C57BL/6^Aire+/-^ (Fig 1F). I. Immune cell flow cytometry of Day 31 post-immunization C57BL/6^Aire-/-^, C57BL/6^Aire+/-^, and C57BL/6^Aire+/+^ mouse prostates.

Histological analysis, scRNA-seq, and cell flow cytometry were performed on C57BL/6^Aire+/+^, C57BL/6^Aire+/-^, and C57BL/6^Aire-/-^ mouse prostates (**Fig 1D**). In C57BL/6^Aire+/+^ and C57BL/6^Aire+/-^ mice, prostatic immune cell populations are minimal (**Fig 1E)**. In contrast, C57BL/6^Aire-/-^ mice consistently develop prostatic inflammation, which persists over time (**Fig 1F**). The immune cell infiltrates are observed most often within the anterior and dorsal prostate lobes, but all lobes are variably affected. Similar to human BPH prostates, immune infiltrates are most notably present within the stroma (**Fig 1F**). These infiltrates are composed predominately of lymphoid cells, which are variably arranged in perivascular and periglandular aggregates to organize lymphoid structures, sometimes forming one or more germinal centers (**Fig 1F**). Lymphoid aggregates are composed of CD4^+^ T cells, CD8^+^ T cells, and CD20^+^ B cells (**Fig.1G, Supplemental Fig.S1**). In the epithelium, plasma cells and lymphocytes often infiltrate the mucosal lamina propria (**Fig 1H**). Additionally, histologically significant inflammation is not observed within select organs (lung, liver, kidney, pancreas) outside of the prostate in all mice, including C57BL/6^Aire-/-^ mice, consistent with previous descriptions of a relatively mild *Aire^-/-^* phenotype in the C57BL/6 background^34,43^.

Consistent with histologic findings, flow cytometry analyses of the prostate immune cell populations show an overall predominance of lymphoid cells with a relatively low proportion of monocyte/macrophage populations (**Fig 1I)**. Additionally, hyperplasia is in both epithelial and stromal cell components is also observed in C57BL/6^Aire-/-^ mice (**Fig 1H)**, consistent with studies demonstrating inflammation-induced epithelial and stromal hyperplasia in BPH.

In all, these results confirm the presence of chronic prostate-confined inflammation within C57BL/6^Aire-/-^ mice but not C57BL/6^Aire+/+^ and C57BL/6^Aire+/-^ mice. The immune cell composition of C57BL/6^Aire-/-^ mouse prostates and human BPH prostates demonstrates several similarities. Additionally, the immune cell composition and morphology in both species, particularly the presence of organizing lymphoid structures, is suggestive of chronic antigenic stimulation and a robust localized adaptive immune response.

### B and T lymphocytes are the dominant immune phenotypes in C57BL/6^Aire-/-^ mouse prostates

To gain further insight into the populations of cells present in all lobes of C57BL/6^Aire-/-^ mouse prostates, scRNA-seq was performed. A total of 14,935 cells were sequenced from pooled prostates of three C57BL/6^Aire-/-^ mice harvested at 21 days post-booster, and 17,149 cells were sequenced from 3 pooled C57BL/6^Aire-/-^ mouse prostates harvested at 35 days post-booster (**Table S2**). The median UMIs detected in each cell was 3,048, and the median number of genes detected per cell was 1,361. Initially, 22 cell clusters were defined (**Fig 2A, Table S3**). Cells in each cluster were classified into various immune and non-immune cell types using canonical marker expression (**Fig 2B, Table S4**) as well as automatic cell type identification using SingleR^42^ (**Fig S2A**). Immune cells are abundant in the scRNA-seq data, with lymphoid cells being the dominant phenotype. Clusters 0, 2, 11, 12, 14, and 20 are B cell clusters (Cd19^+^ Ms4a1^+^ Cd79a^+^ Cd79b^+^). Clusters 1, 3, 4, 9, and 17 are T cell clusters (Cd3d^+^ Cd3e^+^ Cd3g^+^), of which clusters 9 and 17 are NKT cell clusters (Nkg7^+^). T cell clusters 3 and 4 are Cd4^+^ T cells whereas cluster 1 appears to contain cytotoxic CD8^+^ T cells. Clusters 5, 8, and 15 contain fibroblasts (Cola1^+^ Cola2^+^), whereas cluster 7 contains cells that are likely to be smooth muscle cells (Tagln^+^, Acta2^+^). Clusters 10 and 6 are epithelial cells (keratin-expressing cells including Krt5^+^, Krt6a^+^, and Krt18^+^). Clusters 13 and 19 are myeloid cells composed of macrophages/monocytes (Cd68^+^ Itgam^+^) and dendritic cells (Flt3^+^) (**Fig 2B, Table S3**). Cluster 21 contains a small endothelial cell (Pecam1^+^) population. The prevalence of T, B, and myeloid cells observed in this data is reminiscent of that seen in the prostates of men diagnosed with BPH^10,44,45^, though a much greater proportion of B cells are identified in the mouse prostates than in the human BPH specimens.

**Figure 2.**
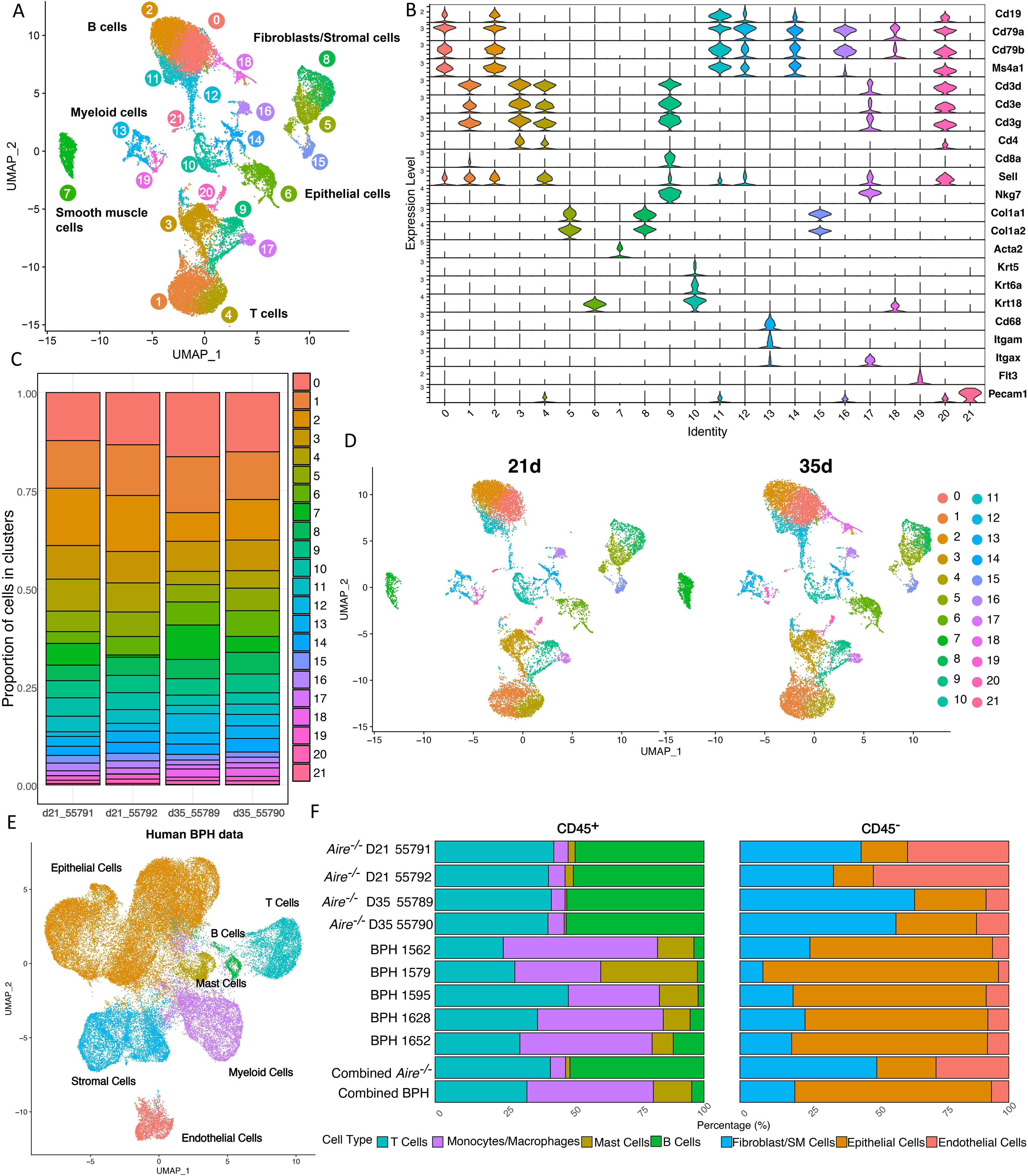
Single-cell RNA-seq of C57BL/6^Aire-/-^ mouse prostates. **A.** Combined clustering of all cells. A total of 22 clusters were detected and were assigned to one of 6 major cell types. **B.** Stacked violin plot of canonical cell markers used to assign cell types to clusters. **C.** A stacked barchart depicts the proportions of cells from each cluster across all Aire^-/-^ samples that were sequenced. **D.** UMAP plot split to separate samples harvested 21 days after receiving a booster or 35 days after receiving a booster. **E.** UMAP of scRNA-seq data from human BPH tissue adapted from^10,14^. **F.** Horizontal stacked barplot depicting the cellular composition of each C57BL/6^Aire-/-^ mouse sample as well as 5 human samples derived from BPH patients^10^. The bottom 2 bars show the cellular composition of combined C57BL/6^Aire-/-^ or human samples.

Overall, there were no significant shifts in the cellular populations or proportions of populations present between prostates harvested 21 days versus 35 days after receiving a booster of prostate homogenate protein and Freund’s Incomplete Adjuvant (**Fig 2C-D**, **S2B**), and there were few statistically significant differentially expressed genes between timepoints (**Tables S5, S6**).

A comparison of the C57BL/6^Aire-/-^ scRNA-seq data to human BPH tissue^10,64^ (**Fig 2E**, **F**) shows considerable similarities and some notable differences between the cellular composition of C57BL/6^Aire-/-^ mouse and human BPH samples. In both C57BL/6^Aire-/-^ and human BPH samples, non-immune cells are dominated by stromal and epithelial cells, with a minor endothelial population of cells present. In C57BL/6^Aire-/-^ mouse, the leukocyte (Cd45^+^) fraction of cells is dominated by lymphocytes whereas in the human BPH samples, lymphocytes and myeloid cell populations comprise a nearly equivalent proportion of the total immune cells. It should be noted, though, that while B lymphocytes are present in human BPH tissue, the proportion of B cells in these data is much lower than in the C57BL/6^Aire-/-^ mouse data. Another notable difference is that mast cells comprise a distinct population of immune cells in the human BPH samples (**Fig 2E**), but in the C57BL/6^Aire-/-^ samples mast cells (*Kit*) do not form a distinct cluster. Furthermore, in BPH patients 3.73% of the total cells are mast cells, whereas we observe fewer mast cells in the prostates of both C57BL/6^Aire-/-^ (1.11% of total cells) and of wild-type C57BL/J6 mice (1.01% of total cells)^44,46^, which were used as a control.

### T cell subclustering indicates the presence of a Gzmk hi T cell subpopulations in C57BL/6^Aire-/-^ prostates

A subclustering of CD3^+^ T cells and CD3^+^ NK cells led to the identification of 14 distinct subclusters (**Fig 3A**). A comparison with young, wild-type C57BL/6J mice^44,46^, which were used as a control, found few T cells present in the prostate (across both samples, only 76 total T lymphocytes were identified; data not shown). The identification of subclusters was both empirically determined using cluster markers (**Fig S3A, Table S7**) and well-known T cell subtype markers (**Fig S3B**) as well as programmatically determined using ProjectTILs^47^ (**Fig 3B, Table S8**). Many populations of T cells show signatures similar to precursor-exhausted T cells (Tpex). An RNA velocity analysis shows that cluster 2 CD8^+^ naïve T cells appear to be an initial cluster, while cluster 7 cells appear to be a terminal cluster (**Fig 3C, D**). Cluster 7 appears to contain a mix of Gzmb hi and low cells, some of which are a Gzmk hi Gzmb low Tox hi subset of Cd8^+^ cytotoxic T cells (**Fig 3E**) that have a similar gene expression pattern to exhausted T cells (**Fig 3B**). The RNA velocity shows that the most terminal population of these cells are Gzmb high, which fits with previous observations regarding granzyme expression and cytotoxic T cell differentiation, specifically that few CD8^+^ T cells express both GZMK and GZMB, and that cells in earlier stages of differentiation tend to be GZMK high GZMB low, and at later stages of differentiation they shift to be GZMK lo GZMB high^48,49^. Interestingly, these Gzmk hi Gzmb low cells appear to be analogous to a subset of CD8^+^ T cells described previously and termed aging-associated T cells (Taa), including the high expression of Cd49d (Itga4)^50^ (**Fig 3E**). An interaction analysis was performed to predict which myeloid cells the Taa cells are interacting with (**Fig 3F**). Cluster 7 Taa cells interact with all 4 macrophage clusters (0,1, 3, and 12), and specifically interacts only with those that show an M1-like or mixed M1 and M2-like phenotype. Notably, the interaction analysis between Taa and myeloid cells mirror those seen in human BPH, in that Taa cells also largely interact with myeloid cells along similar axes (**Fig 3G**).

**Figure 3.**
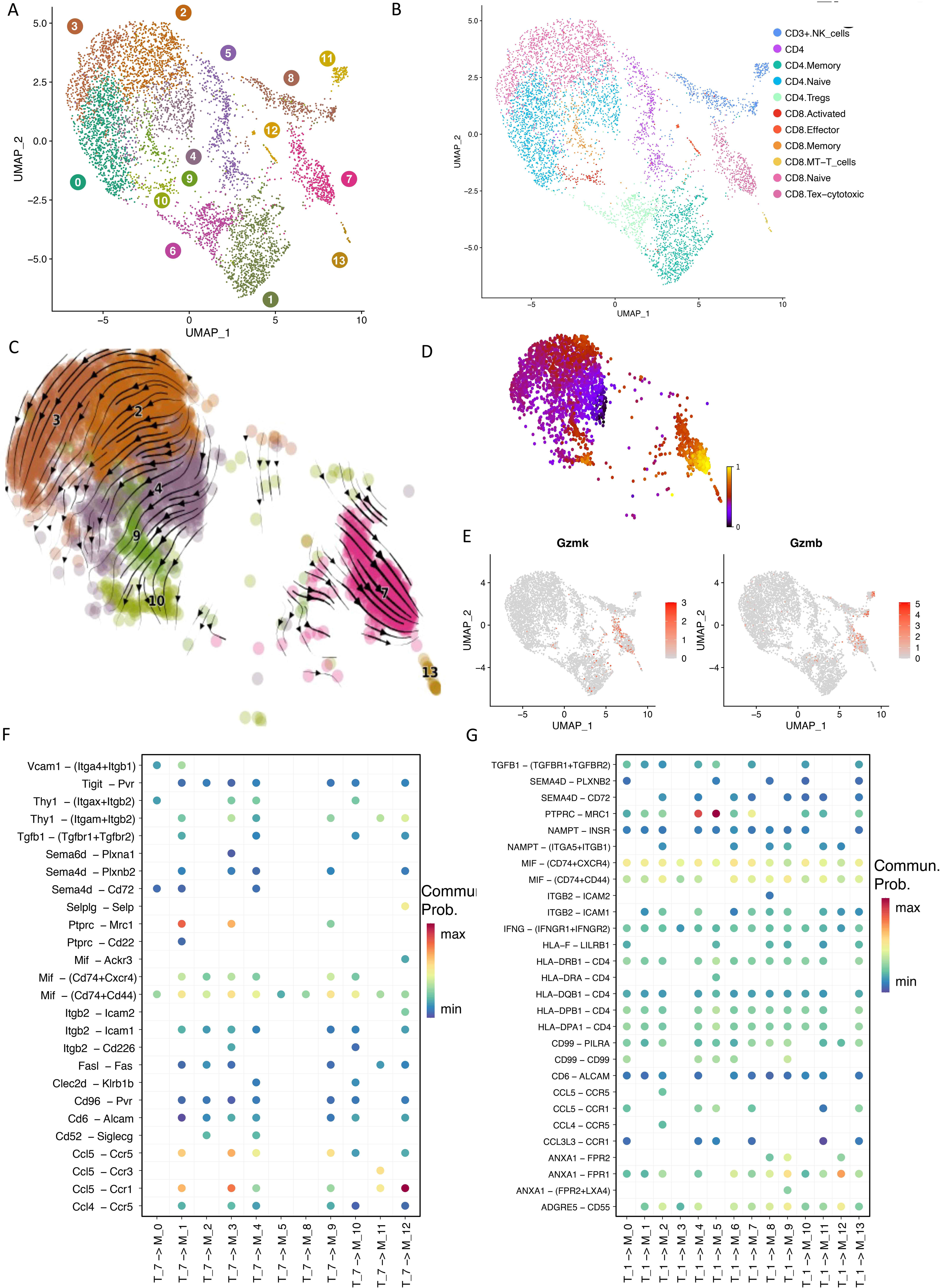
C57BL/6^Aire-/-^ prostate T cell subclustering. **A.** UMAP of the C57BL/6^Aire-/-^ prostate T cell subclustering. **B.** ProjectTILS^47^ annotated UMAP of the C57BL/6^Aire-/-^ prostate T cell subclustering. **C.** Embedding stream diagram of CD8^+^ T cells. **D.** CD8^+^ T cells colored by estimated latent time. **E.** Feature plots highlighting cluster 7 as a mix of CD8^+^ GZMK^+^ TOX^+^ cluster of T cells that is GZMB^-^ and others that are GZMB^+^. **F.** Predicted interactions between aging-associated T cells (T cell cluster 7) and myeloid cell clusters from the C57BL/6^Aire-/-^ prostates. All interactions shown are statistically significant (adjusted p-value<0.01) and points are colored according to the probability of a communication event occurring. The X-axis denotes the clusters that are interacting with T cell clusters expressing the ligands and myeloid cell clusters expressing the receptors. The Y-axis denotes the ligands and receptors involved in the predicted interaction events. **G.** Predicted interactions^72^ between aging-associated T cells (T cell cluster 1) and myeloid cell clusters from prostates of BPH patients^10,14^. All interactions shown are statistically significant (adjusted p-value<0.01) and points are colored according to the probability of a communication event occurring. The X-axis denotes the clusters that are interacting with T cell clusters expressing the ligands and myeloid cell clusters expressing the receptors. The Y-axis denotes the ligands and receptors involved in the predicted interaction events.

### Macrophage subclusters express a spectrum of M1 through M2 polarization signatures in C57BL/6^Aire-/-^ mouse prostates

Myeloid cells are known to increase in numbers in human BPH as prostate size increases, and macrophages have been found to be a highly abundant immune cell population in BPH tissues, second only in number to T cells^10,64^; hence, we performed a subclustering of myeloid cells to investigate the myeloid composition of C57BL/6^Aire-/-^ mice. Overall, 13 clusters of myeloid cells were identified (**Fig 4A**). SingleR^42^ along with marker genes were used to identify clusters of cells (**Fig S4A, 4B, 4C, Table S9**). Clusters 0, 1, 3, and 12 were identified as macrophages (442 cells), clusters 2, 4, 5, 6, 8, and 10 were identified as dendritic cells (473 cells), clusters 7 and 9 were identified as monocytes (117 cells), and cluster 11 as eosinophils (45 cells).

**Figure 4.**
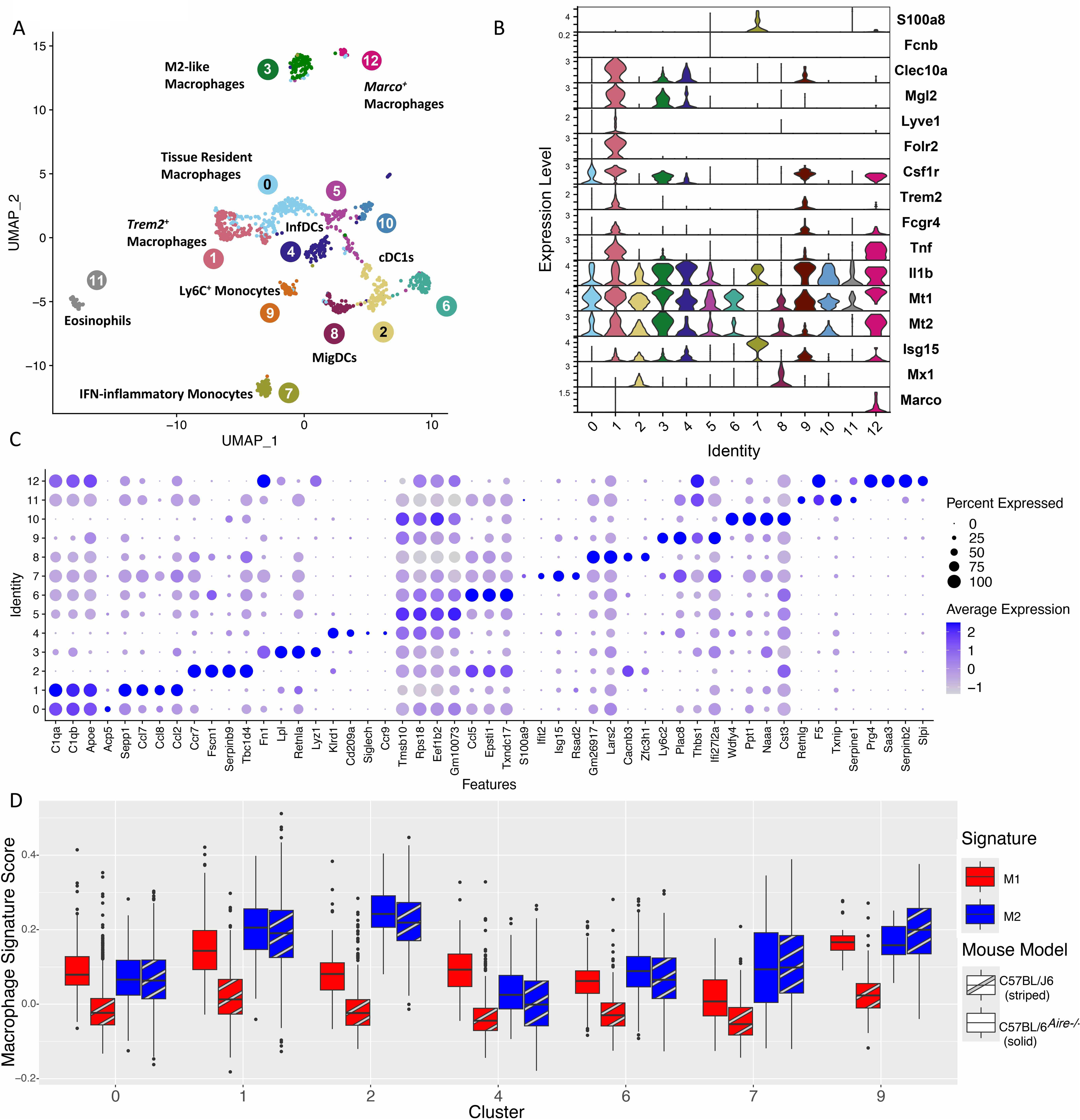
C57BL/6^Aire-/-^ prostate myeloid cell subclustering. **A.** UMAP of the myeloid cell subclustering. A total of 13 distinct clusters were identified and are highlighted. **B.** Stacked violin plot of genes used to aid in myeloid cell identification. **C.** Dot plot showing the statistically significant marker genes (FDR<0.01) of each myeloid subcluster. **D.** Boxplots of the macrophage signature module scores to determine whether an M1 or M2-like gene expression program is expressed by various macrophage populations sequenced from both wild-type C57BL/J6^44^ and C57BL/6^Aire-/-^ prostates.

Amongst the macrophage clusters, cluster 0 and 1 are likely tissue-resident macrophages, as evidenced by the expression of *C1qa*, *C1qb*, and *Apoe*^51^. An analysis to perform supervised classification to identify yolk-sac derived macrophages in the C57BL/6^Aire-/-^ myeloid populations based on a previously identified gene expression signature^52^ further supports that 0 and 1 are likely tissue-resident (**Fig S4C**, **Table S10**). Only cluster 1 has high *Trem2* expression, identified as lipid-associated tissue-resident macrophages that play a potential role in a variety of human conditions^53^, including in BPH^22,54^. *Retnla* is a statistically significant marker of macrophage cluster 3 (**Fig 4C**), suggesting that this is a cluster of alternatively activated (M2) macrophages^55^. Cluster 12 appears to be a population of *Marco*^+^ macrophages (**Fig 4B**), and clusters 1 and 12 also appear to have some inflammatory properties due to the expression of *Tnf* (**Fig 4B**). A recent study found that the abundance of *TREM2*^+^ macrophages and *MARCO*^+^ macrophages is positively correlated with BPH patient body mass index and urinary symptom scores and thus these two populations may play a role in BPH^14^. Also of note, both Cluster 3 and 12 macrophages express high levels of *S100a4*, the gene encoding a calcium-binding enzyme, and similar S100a4^+^ macrophages have been found to increase the development of pulmonary fibrosis in the lung through the activation and proliferation of lung fibroblasts and are secreted by alternatively activated (M2) macrophages^56,57^. Amongst the monocyte clusters, based on the high expression of *Isg15*, *IL1B*, and *S100A8,* it appears that cluster 7 is IFN-inflammatory monocytes (**Fig 4B**, **4C**) that are often found in severe viral infections and autoimmune diseases^58–60^. Cluster 9 (Ly6c2, Thbs1) appears to be a cluster of classical inflammatory monocytes (Ly6C^+^)^61^. Amongst the dendritic cells (DCs), Clusters 10, 2, and 6 (*Xcr1*, *Batf3*, *Irf8*) appear to be conventional type 1 DCs (cDC1s)^62,63^. Cluster 8 (*Cacnb3*) was identified as migratory dendritic cells (migDCs)^64^. DC clusters 4 and 5 (*S100a8*, *Is1b*, *Cd83*) are likely to be inflammatory DCs (infDCs), which have been found to differentiate from monocytes that are recruited to sites of tissue damage and inflammation^65^.

A signature analysis of macrophage subpopulations with M1 and M2 gene expression was performed. This analysis determined that the largest macrophage cluster, cluster 0, has a primarily M1-like phenotype. The signature analysis also determined that cluster 3, not surprisingly given the high expression of *Retnla*, has a predominantly M2-like phenotype, but the other two clusters (1 and 12) appear to have a mixed phenotype (**Fig S4D**)), consistent with TREM2^+^ and MARCO^+^ macrophages in human BPH^14^.

An integrative analysis of the C57BL/6^Aire-/-^ myeloid cells with those of normal wild-type C67BL/J6 mice^44^ was performed and does not identify any clusters of myeloid cells that appear to be unique to the C57BL/6^Aire-/-^ mouse prostate, though the inflammatory dendritic cells in cluster 10 of the integrated clustering are far more prevalent in the C57BL/6^Aire-/-^ mouse prostates (**Fig S4E**, **F**). However, the C57BL/6^Aire-/-^ have a much higher number of M1-like macrophages as well as clusters that appear to have a mixed M1 and M2-like phenotype (**Fig 4D**). The presence of M1-like, M2-like, and mixed phenotypes mirrors what has been seen in macrophage populations derived from the prostates of patients diagnosed with BPH^54^.

### Fibroblasts show the presence of SASP and potential senescence signatures in C57BL/6^Aire-/-^ mouse prostates

Due to the potential that GZMK-expressing Taa cells could contribute to non-resolving inflammation through communication with fibroblasts^66^, a subclustering of all fibroblasts, excluding smooth muscle cells (*Acta2*) was performed (**Fig 5A**) in an attempt to reveal heterogeneity in stromal cells. Fibroblast identity was confirmed for each cluster by both SingleR and marker expression (**Fig S5A**). Fibroblasts formed 12 subclusters in total, and markers were identified (**Table S11**). Fibroblast subclusters were compared with published datasets of fibroblasts from the prostates of young, healthy wild-type mice and human donors^44^ as well as with those from the prostates of BPH patients^44^. Classification of fibroblast subclusters with the gene expression signatures derived from these publicly available datasets using Garnett identifies most cells from cluster 0 (**Fig 5A**, **S4B-E, Table S12**) as being highly similar to human interstitial fibroblasts, such as those identified previously from healthy prostates^44^. Like human interstitial fibroblasts, *Gpx3* and *Apod* are statistically significant (FDR<0.01) markers of cluster 0 (**Table S11**). Clusters 1, 2, 7, and 10 are identified as similar to the prostate-derived fibroblasts from wild-type mice (**Fig 5A**, **S4B, Table S12**) and in total, 91%, 76%, 71%, and 67% of the cells in clusters 1,2, 7, and 10 were classified as wild-type mouse prostate fibroblasts, respectively (**Table S12**). Like the wild-type prostate fibroblasts^44^, clusters 1, 2, and 7 express high levels of the genes *C3*, *Ebf1*, *Gpx3*, *Sult1e1*, and *Igf1* (**Table S11**), which are identified as statistically significant (FDR<0.01) markers of these clusters. Cluster 10 also expresses high levels of these genes. However, *Gpx3* and *Sult1e1* are not statistically significant markers of cluster 10. Clusters 3, 4, 5, and 8 have gene expression profiles similar to two fibroblast populations that were identified from the prostates of men with BPH^44^. They express high levels of *Igfbp2*, a component of the senescence-associated secretory phenotype (SASP) seen in a number of aging-related diseases^67^ and also found at high levels in aged fibroblasts^68,69^. Cluster 6 appears to be the “ductal fibroblasts” described in wild-type mouse prostates (**Fig 5A**, **S4B**, **S4C**, **Table S12**)^44^. The cells in cluster 6 likewise express high levels of *Wnt2*, *Rorb*, *Wif1*, *Ifitm1* and *Srd5a2* (**Fig S4C**, **Table S11**), which were previously identified as markers of ductal fibroblasts in wild-type mice and are statistically significant markers of cluster 6 (FDR<0.01). Clusters 9 and 11 were also annotated ductal fibroblasts classified as ductal (**Fig S4B, Table S12**); however, with the exception of Cluster 11 expressing moderate levels of *Ifitm1*, neither cluster expresses high levels of ductal fibroblast markers (**Fig S4C**, **Table S11**) and none of the ductal fibroblast markers are statistically significant markers of these clusters.

**Figure 5.**
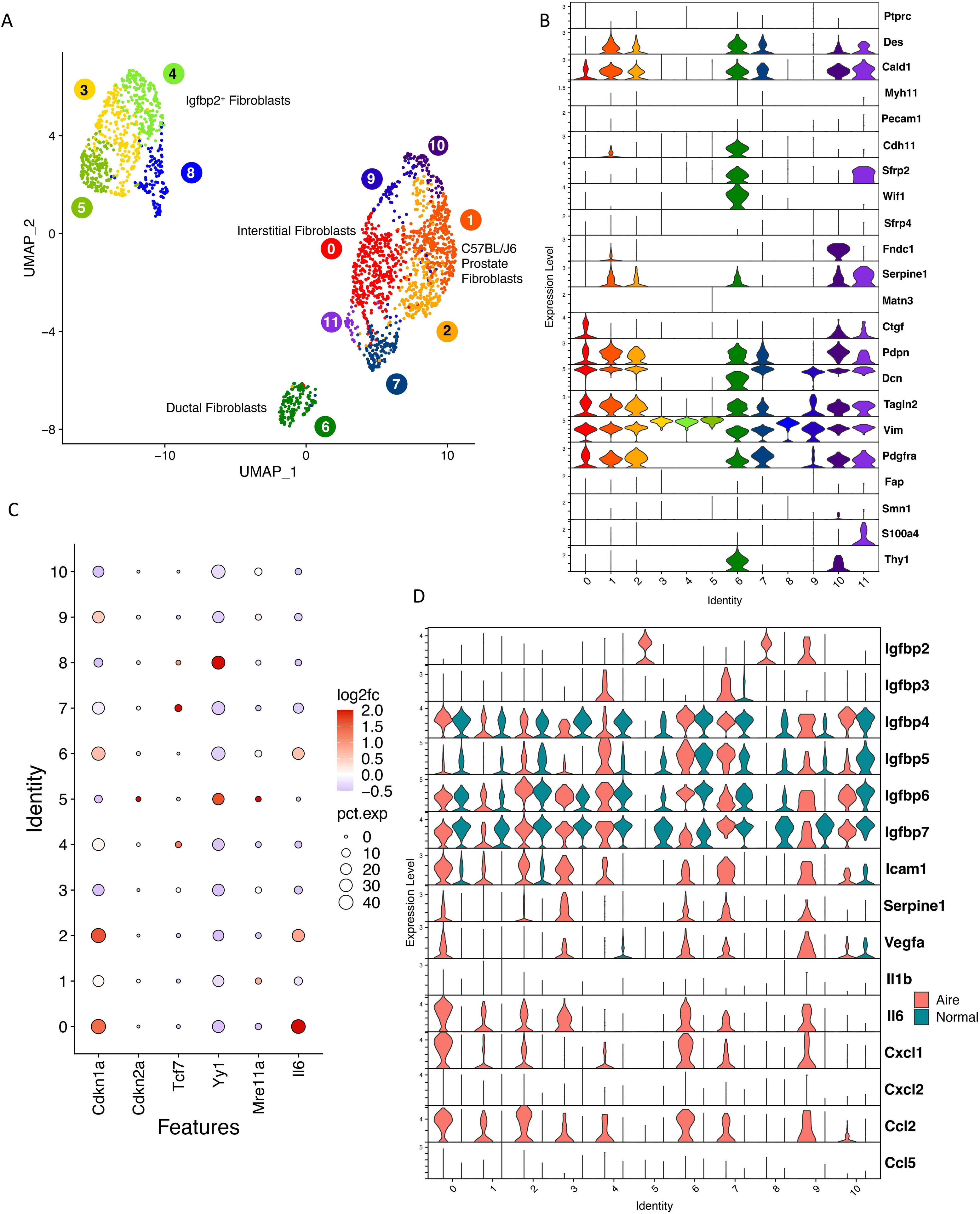
C57BL/6^Aire-/-^ prostate fibroblast subclustering. **A.** Subclustering of fibroblasts. **B.** Senescence-associated gene expression among fibroblast subclusters in C57BL/6^Aire-/-^ mice and in wild-type C57BL/J6 mice^44^, which were combined and integrated. Dots are colored by log_2_fold-change of gene expression in C57BL/6^Aire-/-^ mice compared to wild-type C57BL/J6 mice^44^. **C.** Split stacked violin plot of genes associated with a senescence-associated secretory phenotype (SASP) in C57BL/6^Aire-/-^ mice and in wild-type C57BL/J6 mice^44^.

Fibroblasts derived from the prostates of men diagnosed with BPH also show expression of numerous markers of cellular senescence, specifically of the genes *CDKN1A*, *CDKN2A*, *TCF7*, *ME11A*, *YY1*, and *IL6*^66^. Likewise, a recent study found that fibroblasts from human BPH patients also express high levels of genes associated with a senescence-associated secretory phenotype (SASP)^45^. We next integrated scRNA-seq data from the prostates C57BL/6^Aire-/-^ and normal wild-type C67BL/J6 mice (**Fig S5F**, **S5G**) to determine whether there is a difference in expression of SASP genes. This integrated analysis shows that indeed, the majority of senescence markers show higher expression in C57BL/6^Aire-/-^ than in normal wild-type C67BL/J6 mice (**Fig 5B**). Likewise, all C57BL/6^Aire-/-^ fibroblast clusters express some markers of SASP (**Fig 5C**), and unlike in the wild-type C67BL/J6 mice, the vast majority of clusters in C57BL/6^Aire-/-^ (0,1, 2, 3, 4, 6, 7, and 9) express particularly high levels of such markers.

## Discussion

BPH’s complex factor interplay complicates research model selection. We characterize the C57BL/6^Aire-/-^ inflammation-driven prostatic hyperplasia model, which displays human BPH characteristics including persistent inflammatory macrophage profiles with M1-like macrophages and neither clearly defined M1-nor-M2 phenotypes^54^. Trem2^+^ and Marco^+^ macrophages are present, analogous to T+E2 models^22^ and human BPH prostates where prevalence correlates with symptom severity. C57BL/6^Aire-/-^ prostates exhibit inflammaging markers (SASP, senescence markers in fibroblasts) and Gzmk hi Gzmb low CD8^+^ T cells (aging-associated T cells/Taa) seen in human BPH^45^. Taa cells accumulate with age, contribute to inflammation through Gzmk secretion, making this model useful for studying BPH immune/stromal dysregulation and prostatic inflammaging effects.

*Aire* knock-out C57BL/6 mice injected with prostate homogenate and adjuvant induced localized prostatic inflammation without systemic effects. Harvest timing (21 vs 35 days post-booster) minimally affected cellular composition and gene expression, though limited sample sizes warrant further study.

Notable differences exist between human BPH prostates and C57BL/6^Aire-/-^ mice. Mast cells, abundant in human BPH transition zones and increasing with prostate volume, comprised lower proportions in C57BL/6^Aire-/-^ mice, limiting this model’s mast cell research utility. However, few mast cells in wild-type mice suggest inherent mouse-human differences. While lymphocytes are abundant in both, human BPH samples showed more T than B cells, whereas C57BL/6^Aire-/-^ mice showed the reverse. Human samples also had larger myeloid cell fractions than mouse samples, though human heterogeneity and digestion protocol differences may explain discrepancies.

All mouse models used to study urinary symptoms secondary to BPH have limitations; however, the use of pre-clinical models is essential to unraveling the complex interplay of features that contribute to epithelial and stromal hyperplasia. Future plans include comparing the C57BL/6^Aire-/-^ model with other animal models commonly used to study BPH, including comparisons between immune, epithelial, and stromal cells in the models with cells derived from the prostates of men diagnosed with BPH, as well as comparisons of symptoms such as voiding issues. Such comparisons will be essential for identifying the best pre-clinical models for studying specific aspects of BPH. It is likely that there may be no one “best” model, but that instead, specific models may be useful for studying specific aspects of BPH, such as fibrosis, epithelial and stromal proliferation, and immune cell infiltration. Based on this work, the C57BL/6^Aire-/-^ model is a useful model for studying BPH, particularly the impact of the chronic, non-resolving inflammation in the prostate that is seen in BPH patients. In addition, this model can be used for investigating age-related inflammatory processes (inflammaging) and the contribution of myeloid cell dysfunction to the development and progression of prostate pathology.

## Acknowledgments

The authors are grateful to Dr. Phillip SanMiguel and the Purdue University Genomics Facility for their help sequencing and normalizing the scRNA-seq libraries. This work was also supported by the Collaborative Core for Cancer Bioinformatics (C3B) and the Walther Cancer Foundation. **Funding:** This work was funded through R01-DK126478 (T.L.R.), and P20DK140417 (S.W.H.) from NIDDK, the Purdue University Institute for Cancer Research (P30CA023168), and the IU Simon Cancer Center (P30CA082709).

## Author Contributions

T.L.R. and S.W.H. created the study concept; N.A.L., T.L.R., and S.W.H. designed research studies; M.M.B, D.W., J.Y., A.P.G., B.T.H, S.W.H., and R.E.V. were involved in acquisition of tissues; N.A.L., H.K., and A.K.K. performed statistical and bioinformatics analyses; N.A.L., M.M.B., H.K., A.K.K., J.Y., D.W. and R.E.V. performed data acquisition and all authors contributed to data analysis and interpretation. N.A.L., M.M.B, H.K., T.L.R., S.W.H., and R.E.V. wrote the manuscript and all authors critically revised the manuscript.

---

**Figure S1.**
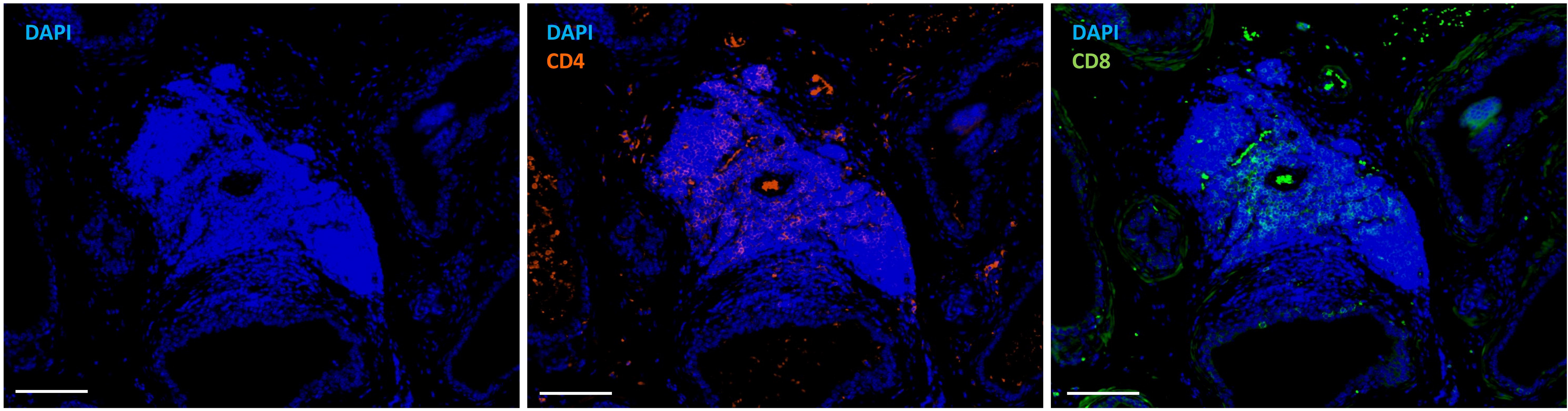
Immunofluorescence of C57BL/6^Aire-/-^ stromal lymphoid aggregate highlighting nuclei (DAPI, left). CD4 (orange, center) or CD8 (green, right) cells. Scale bars=100μm.

**Figure S2.**
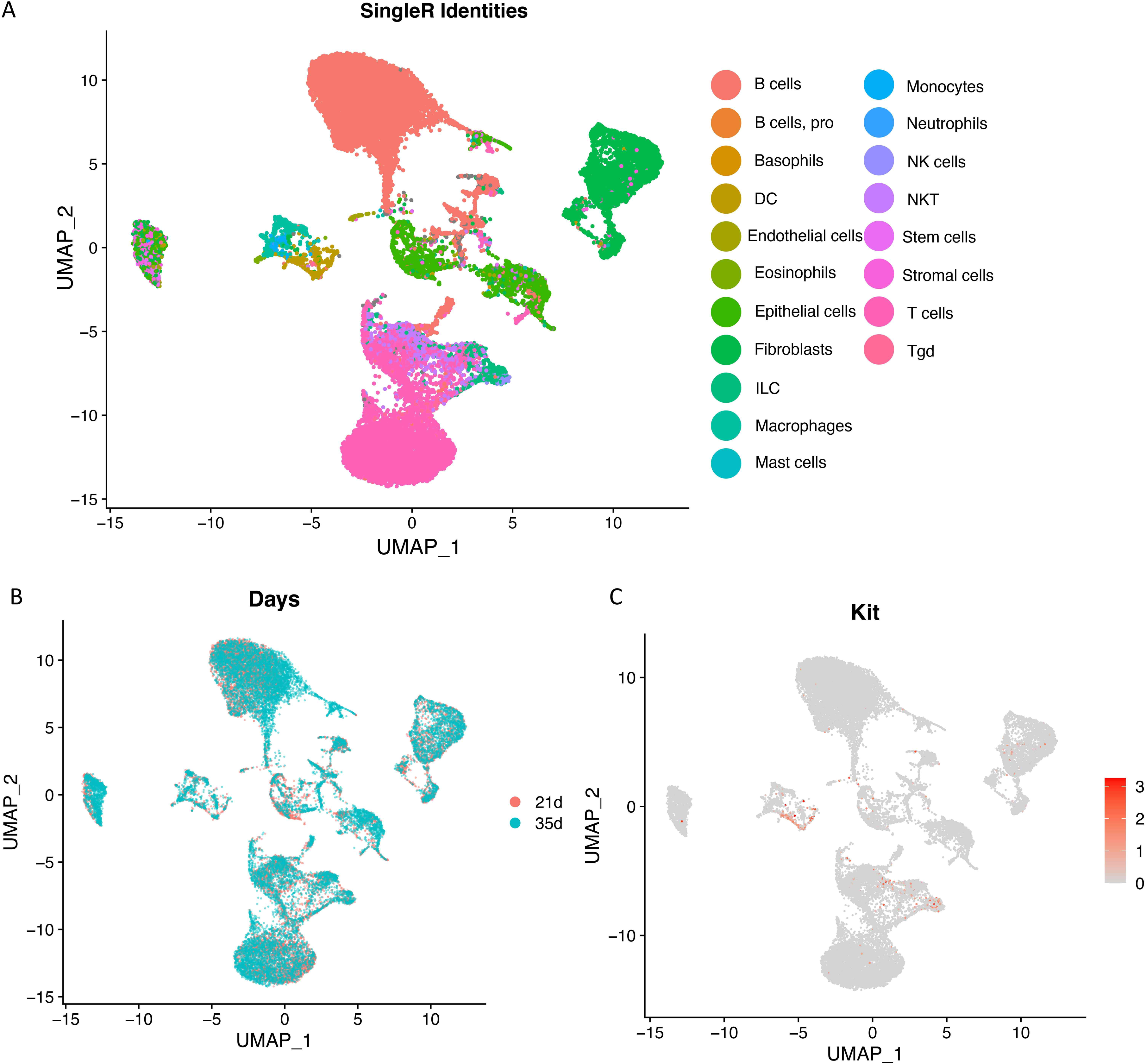
Combined clustering of cells from C57BL/6^Aire-/-^ mouse prostates. **A.** Cell identities as determined by SingleR. **B.** UMAP of C57BL/6^Aire-/-^ mouse prostate-derived cells, colored by the number of days post-booster. **C.** Feature plot showing *Kit* expression, highlighting mast cells.

**Figure S3.**
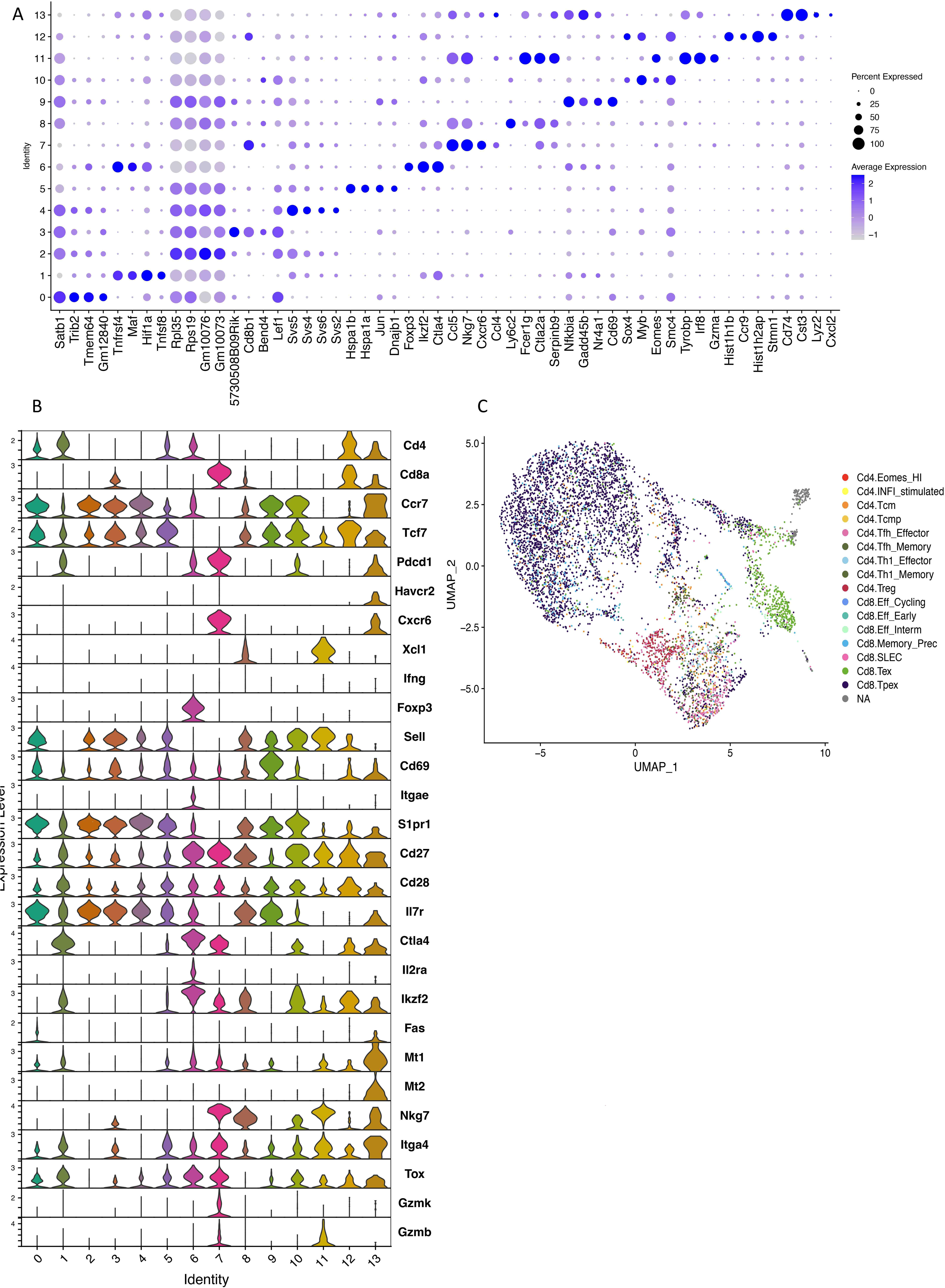
T cell subclustering. **A.** Dotplot of the top 4 statistically significant (FDR<0.01) marker genes identified in each T cell cluster. **B.** Stacked violin plot of genes used to aid annotation of T cell clusters. **C.** ProjectTILS^47^ annotated UMAP.

**Figure S4.**
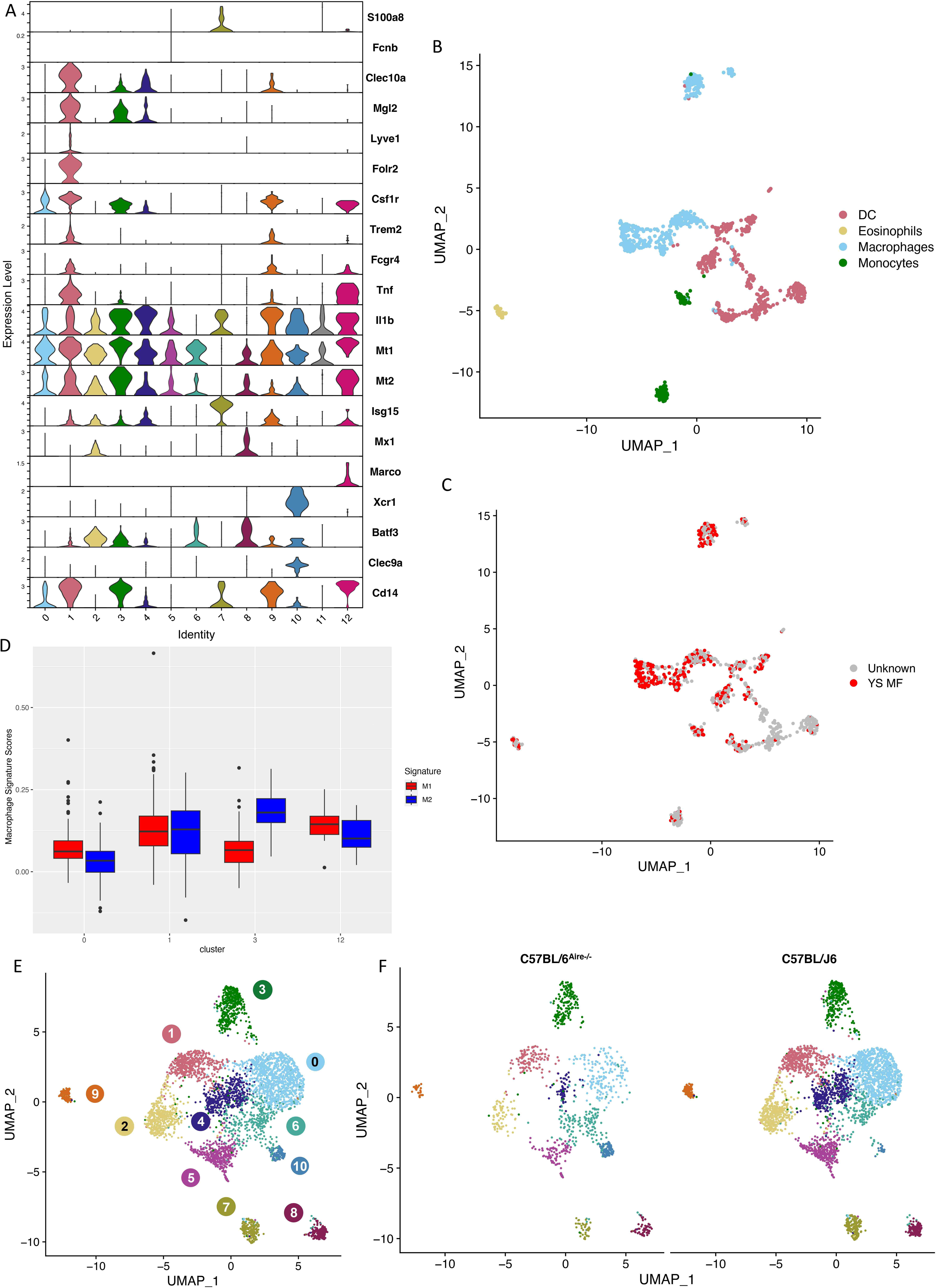
Myeloid cell subclustering. **A.** Stacked violin plot of genes used in annotating myeloid cell populations. **B.** Cluster identities, as determined by the program SingleR. The majority of the cells present are macrophages and dendritic cells. **C.** Supervised classification identifies likely yolk-sac derived macrophages^52^ in the C57BL/6^Aire-/-^ myeloid populations, as shown in the UMAP by red points. **D.** Boxplots of the macrophage signature module scores of the macrophage clusters from the integrated C57BL/6^Aire-/-^ and normal wild-type C67BL/J6^44^ analysis to determine whether an M1 or M2-like gene expression program is expressed by various macrophage populations. **E.** Integrated UMAP showing myeloid cells from C57BL/6^Aire-/-^ and wild-type C57BL/J6 mice^44^. **F.** Split integrated UMAP showing the combined integration of myeloid cells from C57BL/6^Aire-/-^ and wild-type C57BL/J6 mice^44^.

**Figure S5.**
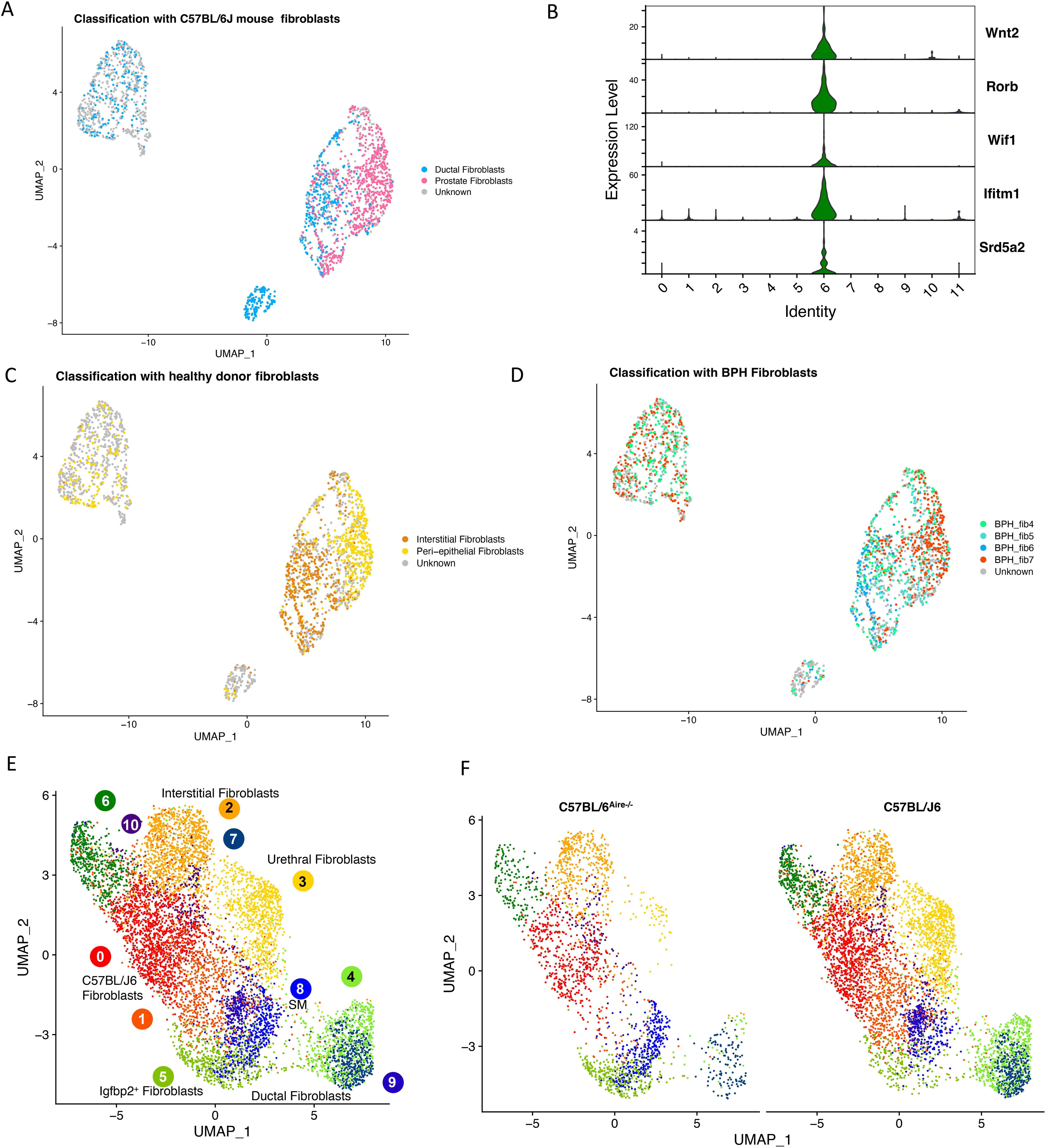
Fibroblast cell subclustering. **A.** Stacked violin plot of marker genes. Gene expression of these markers indicates that these cells are fibroblasts. **B.** Fibroblast subclustering depicting cells classified using markers of prostate-derived fibroblasts from wild-type C57BL/6J mice^44^. **C.** Stacked violin plot of ductal fibroblast marker expression in fibroblast subclustering. **D.** Fibroblast subclustering depicting cells classified using markers of prostate-derived fibroblasts from healthy human prostates. **E.** Fibroblast subclustering depicting cells classified using markers of prostate-derived fibroblasts from the transition zone of men diagnosed with BPH. **F.** UMAP of C57BL/6^Aire-/-^ stromal cells integrated with wild-type C57BL/J6 stromal cells^44^. **G.** Split integrated UMAP showing fibroblasts from C57BL/6^Aire-/-^ and wild-type C57BL/J6 mice^44^.

## Supplemental Tables

**Supplemental Table 2.**
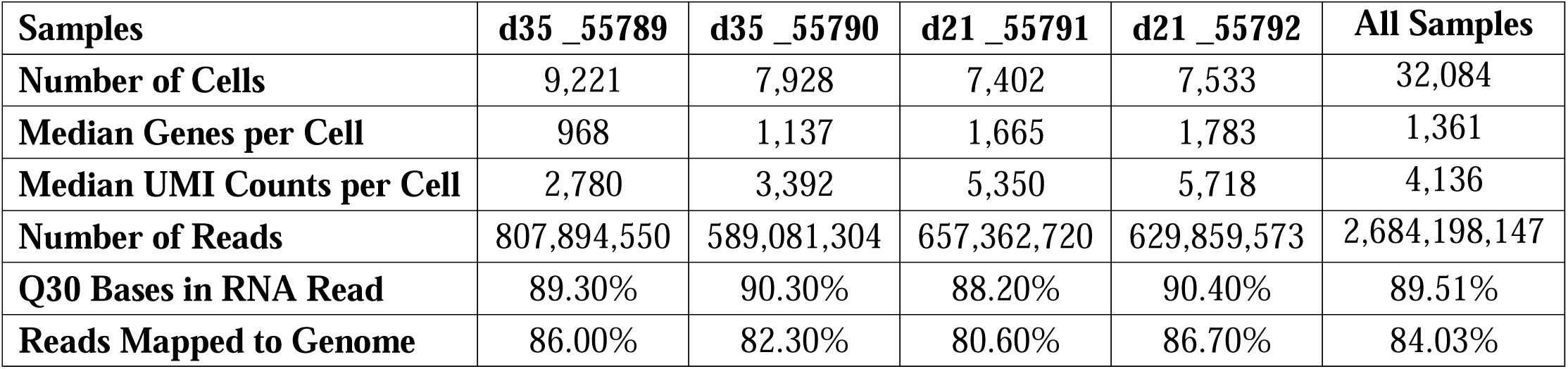
Run Metrics for scRNA-seq of cells from C57BL/6^Aire-/-^ mouse prostates.

**Supplemental Table 3.**
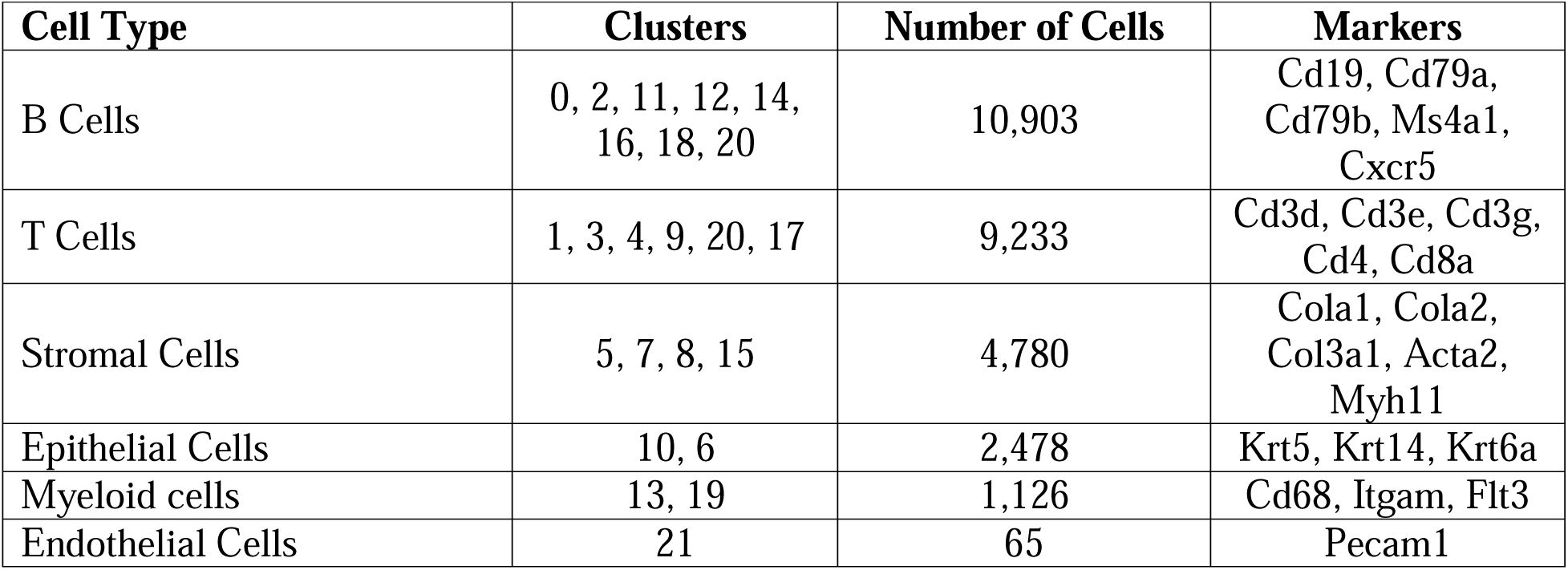
Clusters identified in unsupervised clustering analysis of scRNA-seq data from C57BL/6^Aire-/-^mouse prostates. Cell type and number of cells comprising each cell type are provided as well as the clusters that are associated with that cell type and the marker genes used to identify the cells.

**Supplemental Table 8.**
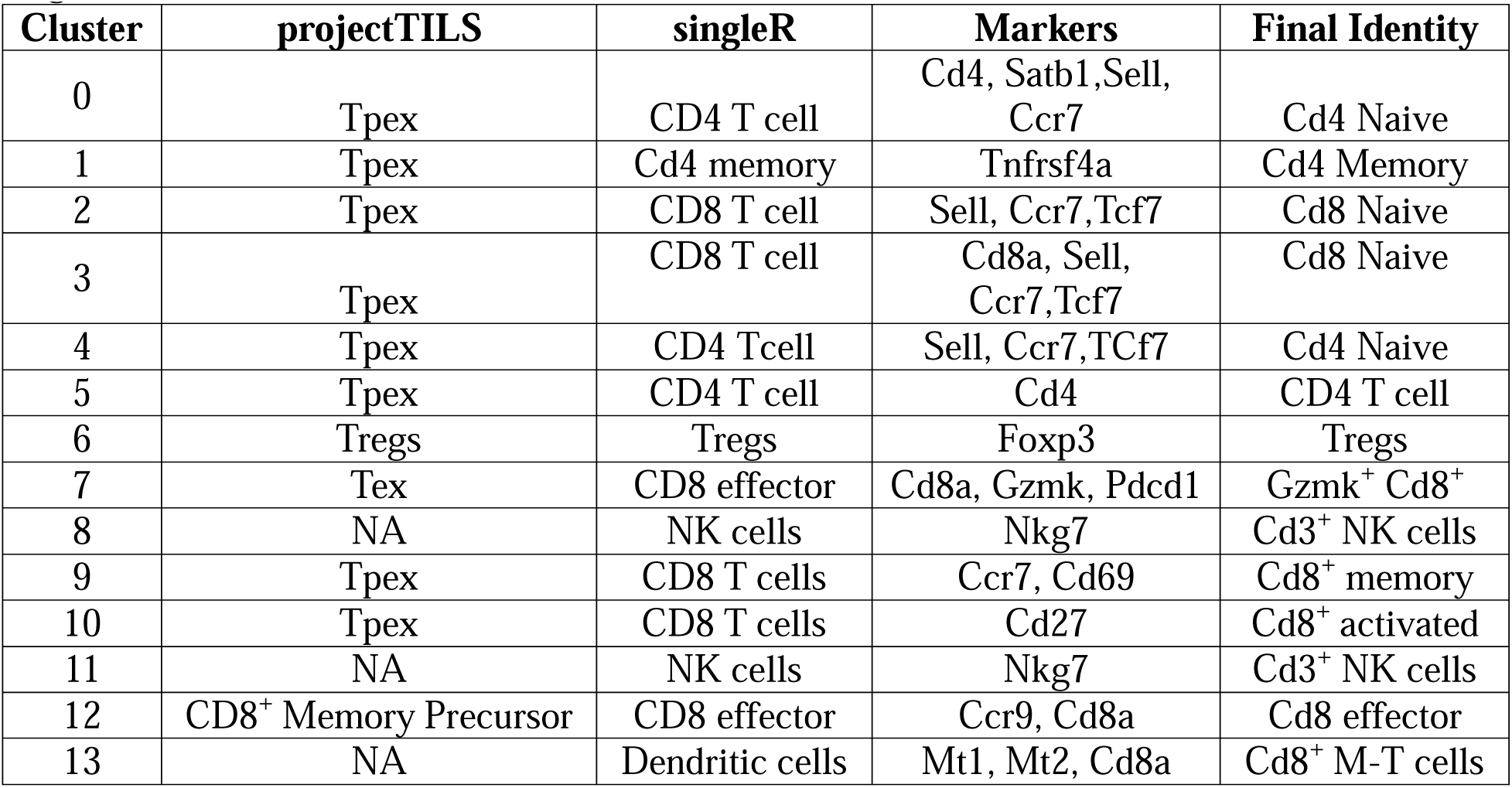
T cell subclustering identities determined by projectTILS, SingleR using the ImmGen database, and the final determination made through marker identification in conjunction with automatic assignment. ProjectTILs and singleR assignments were determined by taking the class of T cells that annotated the majority of cells in a given cluster.

## Table Legends

**Supplemental Table 1. Flow antibody clones.**

**Supplemental Table 2. Run Metrics for scRNA-seq of cells from C57BL/6^Aire-/-^ mouse prostates.**

**Supplemental Table 3. Clusters identified in unsupervised clustering analysis of scRNA-seq data from C57BL/6^Aire-/-^ mouse prostates.** Cell type and number of cells comprising each cell type are provided as well as the clusters that are associated with that cell type and the marker genes used to identify the cells.

**Supplemental Table 4. Marker genes for the combined clustering of all cell types identified in C57BL/6^Aire-/-^ mouse prostates.**

**Supplemental Table 5. Pseudobulk differentially expressed genes identified between timepoints.** Differentially expressed genes were identified across all clusters, as is performed in bulk RNA-seq experiments, between samples harvested at 35 days vs 21 days after receiving a booster. Genes were considered statistically significantly differentially expressed at FDR<5%.

**Supplemental Table 6. Clusterwise differentially expressed genes identified between timepoints.** Differentially expressed genes were identified within each cluster between samples harvested at 35 days vs 21 days after receiving a booster. Genes were considered statistically significantly differentially expressed at FDR<5%.

**Supplemental Table 7. T cell subclustering markers.** Markers were identified using the FindAllMarkers() function in Seurat^40,41^ with an FDR<0.01.

**Supplemental Table 8. T cell subclustering identities determined by projectTILS, SingleR using the ImmGen database, and the final determination made through marker identification in conjunction with automatic assignment.** ProjectTILs and singleR assignments were determined by taking the class of T cells that annotated the majority of cells in a given cluster.

**Supplemental Table 9. Myeloid cell subclustering markers.** Markers were identified using the FindAllMarkers() function in Seurat^40,41^ with an FDR<0.01.

**Supplemental Table 10. Yolk sac-derived macrophages percentages per cluster.** Numbers of cells and percentages of macrophage clusters identified as yolk-sac derived.

**Supplemental Table 11. Fibroblast subclustering markers.** Markers were identified using the FindAllMarkers() function in Seurat^40,41^ with an FDR<0.01.

**Supplemental Table 12. Fibroblast annotation.** Annotations were taken through a combination of marker genes and through the use of marker genes and a multinomial classifier trained based on the top 25 marker genes from fibroblast clusters in BPH^54^, wild-type mice^44^, and healthy human donors^46^, considering 50 unknown cells as the outgroup. The clusters were assigned a particular annotation based on the consensus of cell annotations within that cluster. This annotation was only adopted if more than 50% of the cells within that cluster shared the same annotation, and the cluster’s marker genes were consistent with those known for the reference cell population.

## Notes

### Competing Interest Statement

The authors have declared no competing interest.

